# Kinesin KIF18A is a novel PUM regulated target promoting mitotic progression and survival of human male germ cell line - TCam-2

**DOI:** 10.1101/819839

**Authors:** Maciej Jerzy Smialek, Bogna Kuczynska, Erkut Ilaslan, Damian Mikolaj Janecki, Marcin Piotr Sajek, Kamila Kusz-Zamelczyk, Jadwiga Jaruzelska

**Affiliations:** Institute of Human Genetics, Polish Academy of Sciences, Poznan, Poland; Institute of Bioorganic Chemistry, Polish Academy of Sciences, Poznan, Poland

## Abstract

Regulation of proliferation, apoptosis and cell cycle is crucial for the physiology of germ cells. Their malfunction contributes to infertility and germ cell tumours. Kinesin KIF18A is an important germ cell specific regulator which downregulates apoptosis while promoting cell proliferation in animal models. Whereas regulation of KIF18A expression was studied at the transcriptional level, its posttranscriptional regulation has not been extensively explored. Due to the presence of two PUM Binding Elements (PBEs) within 3’UTR, *KIF18A* mRNA is a potential target of PUMs, well known RNA-binding proteins involved in posttranscriptional gene regulation (PTGR). We investigated that possibility in TCam-2 cells originating from seminoma, representing human male germ cells. We conducted RNA co-immunoprecipitation combined with RT-qPCR, as well as luciferase reporter assay by applying appropriate luciferase construct encoding the wild type *KIF18A* 3’UTR, upon PUM1 and PUM2 overexpression or knockdown. We found that KIF18A is repressed by PUM1 and PUM2. To study how this regulation influences KIF18A function in TCam-2 cells, MTS proliferation assay, apoptosis and cell cycle, analysis using flow cytometry was performed upon *KIF18A* siRNA knockdown. We uncovered that KIF18A significantly influences proliferation, apoptosis and cell cycle, these effects being opposite to PUM effects in TCam-2 cells. We propose that repression by PUM proteins may represent one of mechanisms influencing KIF18A level in controlling proliferation, cell cycle and apoptosis in TCam-2 cells. To the best of our knowledge, this paper identifies the first mammalian functionally germ cell specific gene that is regulated by Pum proteins via 3’UTR.

## INTRODUCTION

Infertility is a worldwide problem which affects about 15% of couples globally, half of them being caused by the male factor (for review see Inhorn and Patrizio, 2015). Despite the availability of various medical procedures, such as artificial reproductive technology, healing of infertility is not possible in many cases (Bhartiya, 2015). It is believed that dysfunction of fertility may occur during fetal life of an individual, at the time when the germline is established, or may take place later, during the germ cell maturation. It is of note that male infertility is frequently associated with testicular germ cell tumours, thus considered as a risk factor (Matzuk and Lamb, 2008). Kinesins are enzymatic proteins playing therole of molecular motors in intracellular transportation of cargo along microtubules and in cellular movements (Sperry, 2012), these activities being catalysed by ATP hydrolysis (Vale and Milligan, 2000). Depletion of kinesin-8 family of proteins causes abnormalities in mitotic spindle lengthening (Goshima et al., 2005), chromosome alignment failure (Stumpff et al., 2008) and cell cycle dysfunction (Straight et al., 1998). KIF18A supports chromosome congression during mitosis by reducing the amplitude of kinetochore oscillations in pre-anaphase and slows down poleward movements in anaphase (Stumpff et al., 2008).

Although Kif18a is ubiquitously expressed, its role in mice was demonstrated to be limited to germ cell development (Czechanski et al., 2015). Namely, point mutation of the Kif18a motor domain in germ cell lineage in fetal gonads leads to chromosome alignment defects and mitotic arrest (in G2/M phase), due to spindle assembly MAD2 checkpoint activation. (MAD2 localisation to kinetochores provides a signal that inhibits progression from metaphase to anaphase) (Li and Benezra, 1996). Those abnormalities trigger apoptosis resulting in germ cell depletion and infertility (Czechanski et al., 2015). Interestingly, Kif18a dysregulation in somatic cells, despite chromosome alignment defects, does not cause neither mitotic arrest nor apoptosis (Czechanski et al., 2015). Such discrepancy confirms the unique function of Kif18a in germline development. Although KIF18A orthologue was demonstrated to be involved in the mouse spermatogenesis, its precise role in human germ cells has not been studied yet.

Notably, the expression of kinesins-8 is often dysregulated in cancer and therefore may play an important role in that process. Interestingly, overexpression of KIF18A was reported to promote tumour formation, invasion and metastasis in colorectal cancer, while its knockdown triggered proliferation decrease in cancer cell lines, altogether indicating a potential oncogenic activity (Nagahara et al., 2011). Upregulation of KIF18A in various types of human cancers indicates that accurate regulation of KIF18A level is pivotal for preserving proper cell cycle progression. Although there is a report on transcriptional regulation of KIF18A expression potentially mediated by FOXM1 transcriptional factor (Muller et al., 2014), as well as posttranscriptional regulation in cancer via TTP and HUR RNA-binding proteins (Hitti et al., 2016), the mechanisms of KIF18A regulation were not studied extensively.

PUM proteins are highly conserved RNA-binding proteins that target mRNAs by binding 8-nt long consensus PUM Binding Element (PBE) UGUAHAUW, located mostly within 3’UTR (Bohn et al., 2018). PBE recognition by PUMs is mediated by a highly conserved PUF-domain and depends on PUM cooperation with several protein cofactors (Bohn et al., 2018; Wickens et al., 2002) (Smialek et al. [PDF] biorxiv.org, doi: https://doi.org/10.1101/760967). Both mammalian paralogues, *PUM1* and *PUM2*, are very similar in structure (Spassov and Jurecic, 2003), and PUM1 was shown to be important for the mouse germline development (Chen et al., 2012). We sought to investigate its potential regulation by PUM proteins due to presence of two PBEs within its 3’UTR which we selected in our recent screens for PUM1 and PUM2 targets in TCam-2 cells (Smialek et al. [PDF] biorxiv.org, doi: https://doi.org/10.1101/760967). These cells originate from human seminoma, a type of testis germ cell tumour (TGCT) representing human male germ cells at an early stage of prenatal development (de Jong et al., 2008). Notably, these two PBE (UGUAHAUW) motifs are conserved in mammals, given that both are present in Kif18A 3’UTR of the mouse homologue (not shown). Our results indeed show that KIF18A is regulated by PUM1 and PUM2 proteins, they both repress KIF18A expression and this repression may impact the effect of KIF18A on proliferation, apoptosis and the cell cycle progression of TCam-2 cells.

## RESULTS

### KIF18A expression is repressed by PUM1 and PUM2 proteins in TCam-2 cells

We have previously selected *KIF18A* mRNA as being a candidate target of PUM1 and PUM2 in posttranscriptional gene regulation (PTGR) in TCam-2 cells by combining CLIP-Seq and RNA-Seq (Smialek et al. [PDF] biorxiv.org, doi: https://doi.org/10.1101/760967). The *KIF18A* mRNA contains two PBE motifs in 3’UTR (**Fig.1A**). To confirm our preliminary results, we have performed overexpression (OE) of PUM1 or PUM2 in fusion with DDK tag in TCam-2 cells, followed by UV crosslinking and RNA co-immunoprecipitation using anti-DDK antibody. By this approach we have shown binding of *KIF18A* mRNA to PUM1 and PUM2 (**Fig.1B**). Furthermore, we have demonstrated *KIF18A* mRNA and protein upregulation upon an efficient *PUM1* and *PUM2* knockdown, in comparison to cells treated with control siRNA (**Fig.1C & 1D**). By these two approaches we demonstrated that KIF18A is under PUM1 and PUM2 repression.

**Fig 1.**
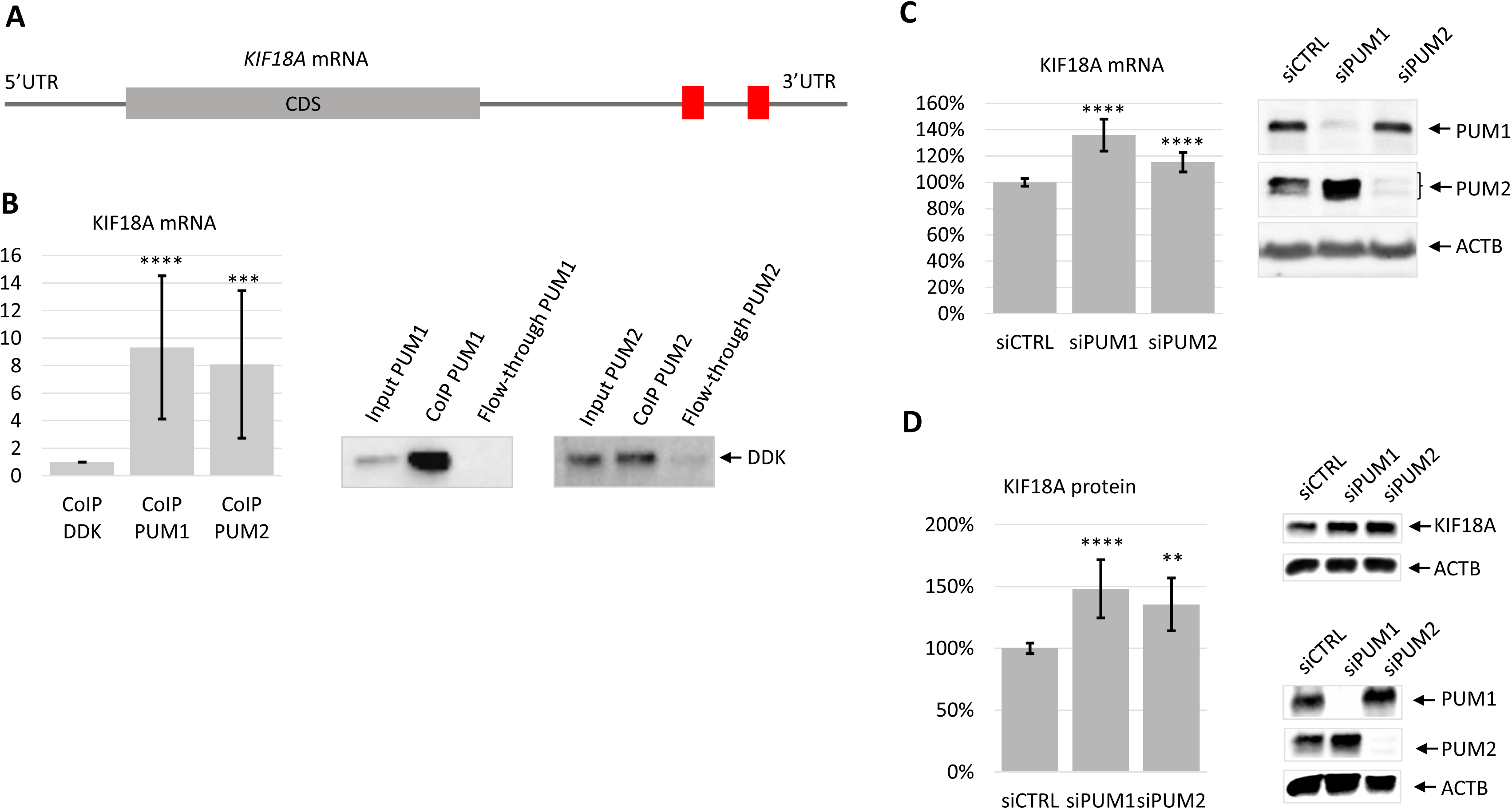
The influence of PUM1 and PUM2 proteins on KIF18A expression. **A** - Scheme of *KIF18A* mRNA with PBE motifs (UGUAHAUW) within 3’UTR represented as red boxes. **B** - RT-qPCR showing enrichment of *KIF18A* mRNA in PUM1 and PUM2 RIP obtained using anti-DDK antibody compared to pCMV6-entry empty vector as a negative control; Additionally *Act5c* mRNA from *Drosophila melanogaster* spike-RNA was measured serving as a reference gene for RT-qpCR (left panel). Representative western blot showing PUM1 and PUM2 binding to beads (right panel). This experiment was performed in 3 biological replicates and 5 technical repeats in each. **C** - RT-qPCR showing *KIF18A* mRNA upregulation upon PUM1 and PUM2 siRNA silencing, as measured 72h post-transfection. *ACTB* and *GAPDH* mRNAs were used as reference genes for RT-qPCR (left panel). Representative western-blot showing efficiency of PUM1 and PUM2 silencing are shown (right panel). Experiments were performed in 3 biological replicates and 5 technical repeats in each. **D** - Upregulation of KIF18A protein level upon PUM1 and PUM2 silencing. Representative western-blots showing KIF18A protein upregulation (upper right panel), as an effect of PUM1 and PUM2 protein silencing (lower right panel). ACTB served as reference in western blot experiments. The experiment was performed in 3 biological replicates and 3 technical repeats. ** P-value<0.005; *** P-value<0.0005; **** P-value<0.00005

As the next step, we sought to investigate whether regulation by PUM proteins is mediated by *KIF18A* 3’UTR containing two PBE motifs. For that purpose we prepared a construct encoding the full-length *KIF18A* 3’UTR downstream of Renilla luciferase (**Fig.2A**). We performed dual-luciferase assay to measure expression of that construct 24 h post PUM1 and PUM2 overexpression, compared to empty pCMV6-entry vector as a negative control. We have shown that upon PUM1 and PUM2 overexpression *KIF18A* 3’UTR was downregulated (**Fig.2B** left panel), in comparison to the empty pCMV6-entry vector overexpression. A significantly weaker effect of PUM2 compared to PUM1 could be accounted for by a much lower PUM2 overexpression level compared to PUM1 (**Fig. 2B** right panel). To confirm that result, we performed *PUM1* and *PUM2* siRNA knockdown, in comparison to control siRNA. As expected, we observed upregulation of luciferase construct expression (**Fig.2C** left panel) upon an efficient *PUM1* and *PUM2* knockdown (**Fig.2C** right panel). Both approaches show that *KIF18A* mRNA is repressed by PUM1 and PUM2 in TCam-2 cells, and this effect is mediated by 3’UTR.

**Fig. 2.**
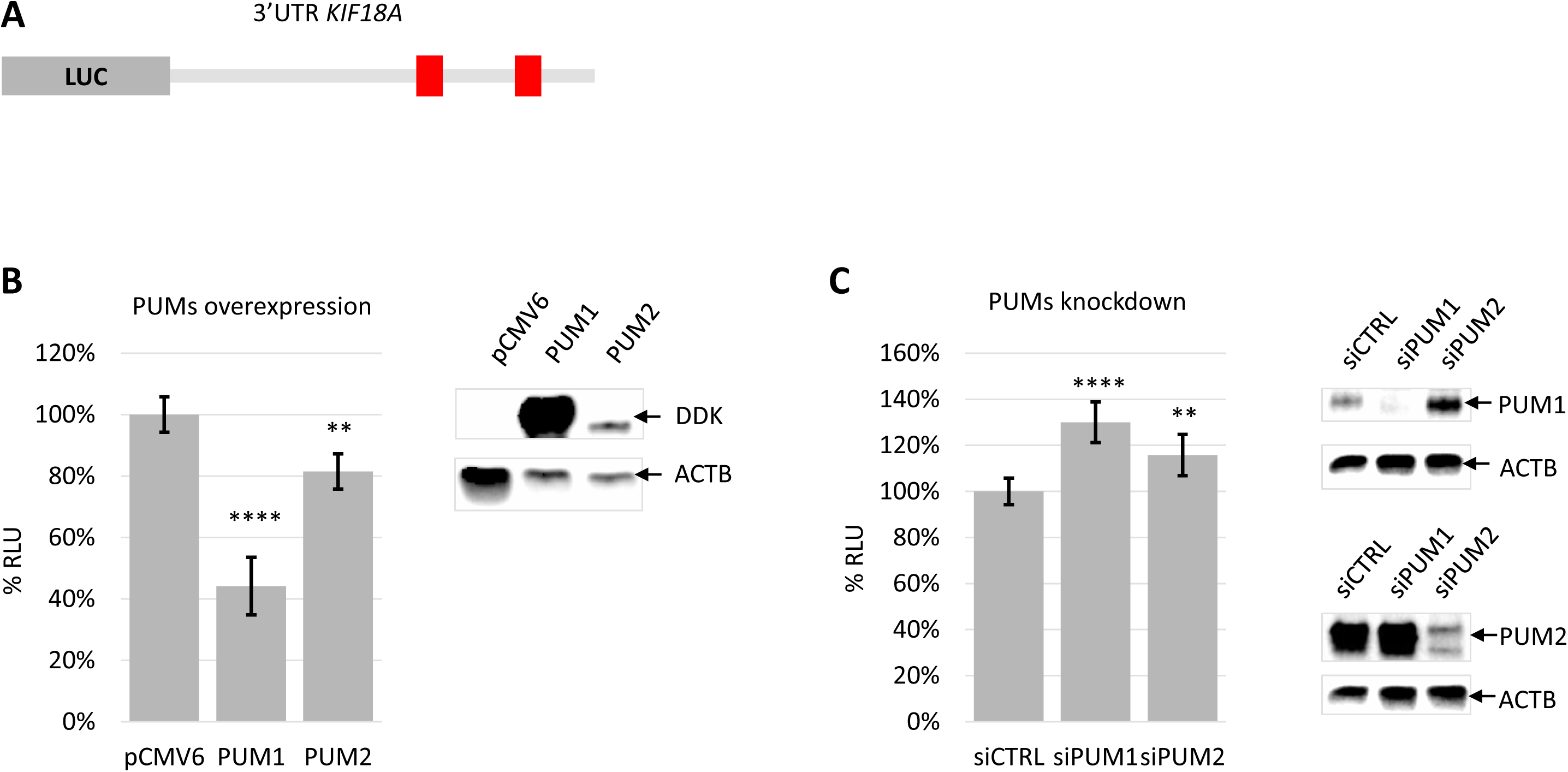
The effect of PUM1 and PUM2 on the expression of *KIF18A* LUC-3’UTR construct measured by dual-reporter luciferase assay. **A** - The scheme of *KIF18A* LUC-3’UTR construct with red boxes indicating Pumilio Binding Elements (PBEs – UGUGAHAUW). **B** - The effect of PUM1 and PUM2 overexpression on the expression of *KIF18A* LUC-3’UTR construct. Luminescence values (RLU = relative luciferase units) are expressed as % of the RLU of samples transfected with reporter construct only, which was set to 100%. *Renilla* luciferase values were normalised using firefly luciferase measurements (left panel). Representative western-blot showing PUM1 and PUM2 overexpression level (anti-DDK antibody); ACTB serving as reference (anti-ACTB antibody) (right panel). **C** - The effect of *PUM1* and *PUM2* knockdown on *KIF18A* LUC-3’UTR construct expression (left panel). The level of construct up-regulation upon PUMs silencing is shown as % RLU in comparison to control siRNA set to 100%. Representative western-blot showing the level of PUM1 and PUM2 silencing, ACTB serving as reference (right panel). All experiments were performed in 3 biological replicates with 2 technical in each. ** P-value<0.005; **** P-value<0.00005

### Knockdown of KIF18A caused decreased TCam-2 cell proliferation, while PUMs’ knockdown induced opposite effect

Since it was reported that Kif18a influences mouse germ cell proliferation (Czechanski et al., 2015; Luo et al., 2018; Zhong et al., 2019), we first investigated such potential effect of human KIF18A on TCam-2 cells, which originates from male germ cells. For that purpose we performed MTS proliferation assay upon *KIF18A* knockdown, starting at 24 h up to 120 h post transfection. We observed that proliferation of cells decreased upon *KIF18A* knockdown and this effect was observed during that period, in comparison to cells with control knockdown (siCTRL).

To investigate the influence of PUM proteins on proliferation, we performed MTS proliferation assay upon *PUM1* and *PUM2* siRNA silencing. We found that both *PUMs’* knockdowns resulted in upregulation of cell proliferation in comparison to cells with control knockdown (siCTRL). Moreover, positive effect of PUM1 knockdown on cell proliferation was more pronounced than PUM2. This effect of *PUMs*’ knockdown was opposite to *KIF18A* knockdown (**Fig.3A**). The expression of *PUM1, PUM2* and *KIF18A* was efficiently silenced as shown in **Fig.3B**, starting from 48 h post-transfection time-point.

**Fig. 3.**
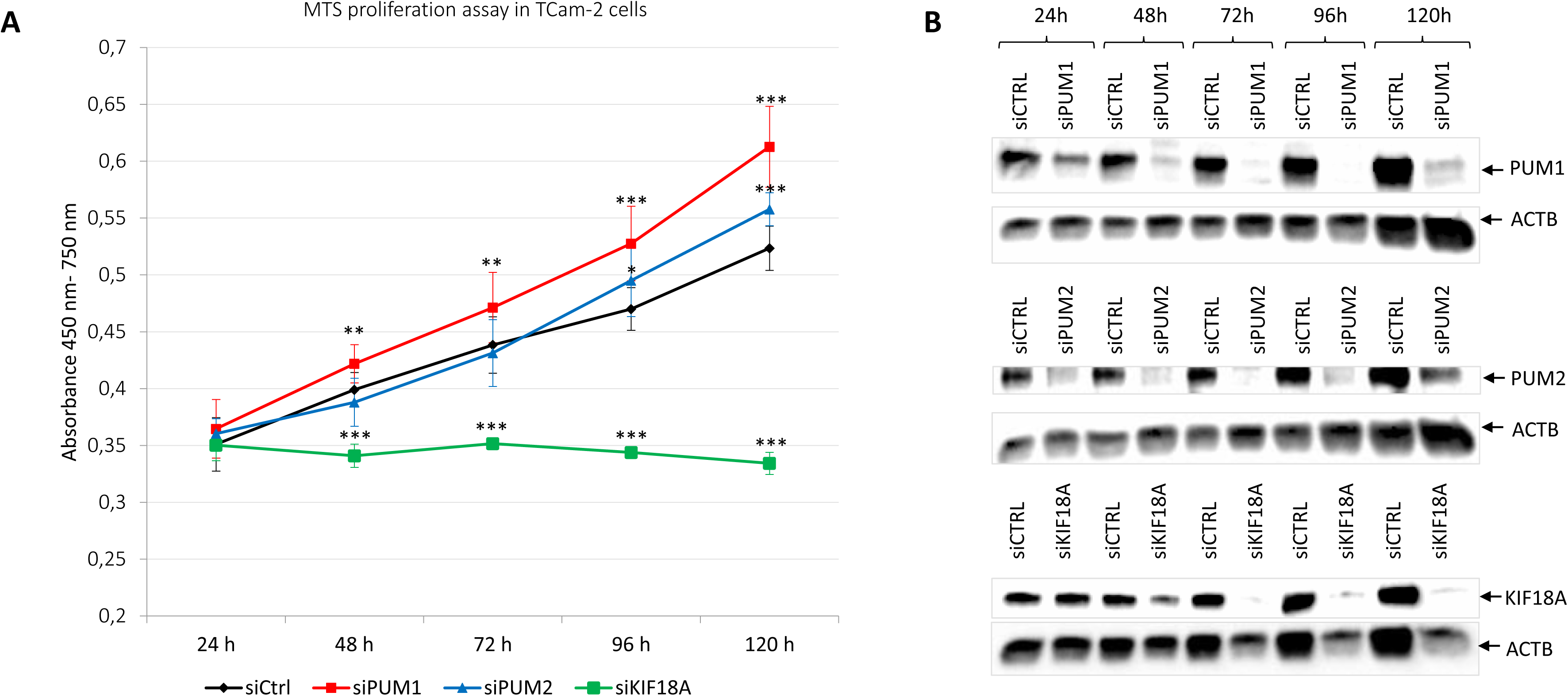
Effect of KIF18A, PUM1 and PUM2 knockdown on cells proliferation. **A** - The MTS proliferation assay was performed on cells 24, 48, 72, 96 and 120 h post-transfection with KIF18A, PUM1 or PUM2 siRNA (each point represents an average of 3 biological replicates (with 7 technical in each)); absorption measurements were performed at 450nm (Formazan) minus 750nm (background) 4h after addition of MTS reagent. **B** - The representative western-blots showing efficiency of KIF18A siRNA or PUMs silencing starting from 24 h post-transfection in comparison to ACTB serving as reference. * P-value<0.05; ** P-value<0.005; *** P-value<0.0005

### KIF18A knockdown causes significant upregulation of apoptosis in TCam-2 cells which is opposite to PUM proteins effect

It was shown that depletion or a single amino acid substitution in Kif18a caused elevated apoptosis exclusively in germ cells in mice, with no effect on somatic cells (Czechanski et al., 2015). Also, *Kif18a* knockout stimulated apoptosis, which resulted in the absence of germ cells in seminiferous tubules and infertility (Czechanski et al., 2015). Therefore, we investigated potential influence of human *KIF18A* on apoptosis in TCam-2 cells. We have observed that upon *KIF18A* knockdown population of TCam-2 cells was reduced by half (data not shown). In another half of cells which survived, almost 2.5 times increase in apoptosis ensued, in comparison to control cells (**Fig. 4A** upper panel). The knockdown efficiency of *KIF18A* is shown in **Fig.4A** lower panel. KIF18A depletion caused significant increase in apoptosis in TCam-2 human germ cell model. The effect on apoptosis is visible on representative dotplots (**Fig.4B** left and right panel).

**Fig. 4.**
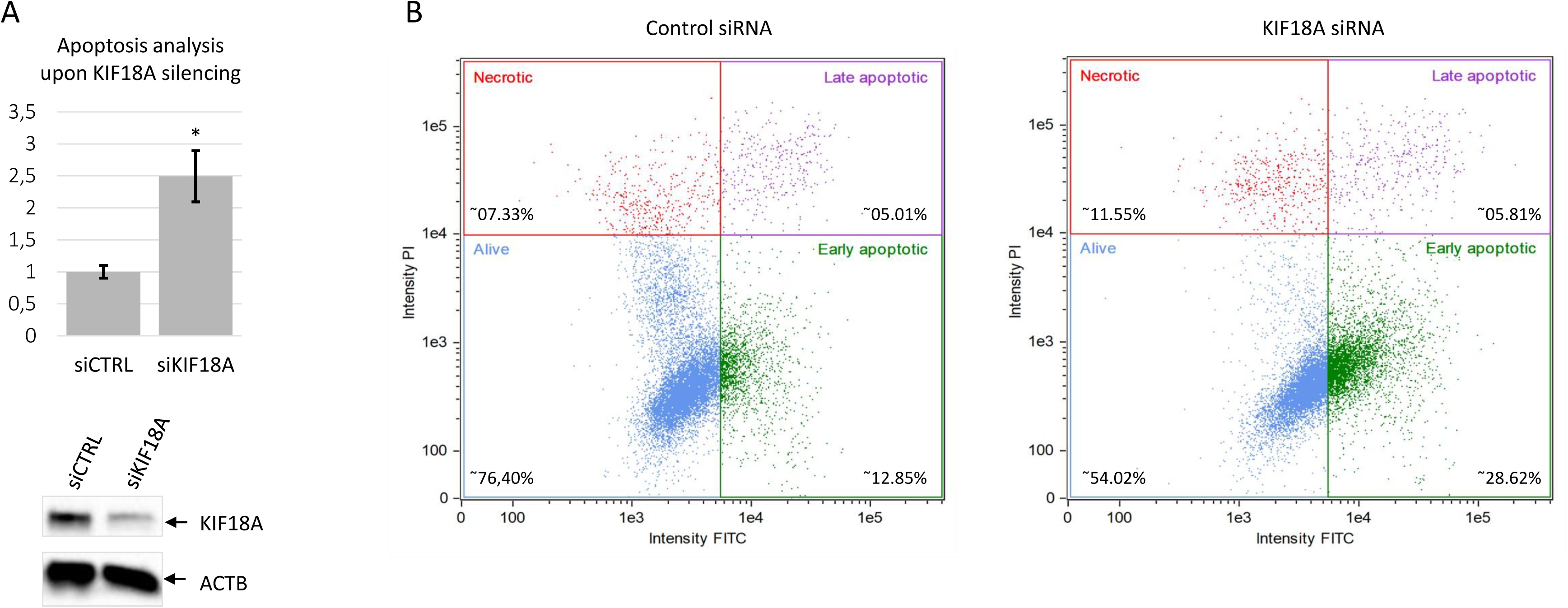
The effect of KIF18A siRNA silencing on apoptosis, as measured using imaging flow cytometry and Annexin V staining assay. **A** - Graph showing the effect of KIF18A knockdown on apoptosis in comparison to control apoptosis level set as 1 (upper panel). Representative western-blot using anti-KIF18A antibody showing efficiency of KIF18A silencing 48h post-transfection in comparison to control siRNA (siCTRL). ACTB detection using anti-ACTB antibody served as the reference (lower panel). **B** - Representative Amnis Ideas graphs showing alive, early apoptotic, late apoptotic and necrotic cells 48h post-transfection with control (left panel) and KIF18A siRNA (right panel). All experiments were performed in 3 biological replicates with 2 technical in each. * P-value<0.05

Given the effect of KIF18A on apoptosis, as the next step we sought to identify apoptotic genes regulated by KIF18A in TCam-2 cells. To this aim, upon *KIF18A* siRNA knockdown in TCam-2 cells, 48 and 72 h post-transfection we performed RT-qPCR measurement of five anti-apoptotic genes: *AKT1, BCL2, CREB1, CCND1* and *PIK3CA* and one pro-apoptotic *BCL2L1* (shorter isoform). All of them are highly expressed in TCam-2 in our previous RNA-Seq screens (Smialek et al. [PDF] biorxiv.org, doi: https://doi.org/10.1101/760967). Our analysis showed that KIF18A knockdown caused a significant downregulation of anti-apoptotic *BCL2* mRNA level, and upregulation of anti-apoptotic *CCND1* and *PIK3CA*. Upregulation of pro-apoptotic shorter isoform of *BCL2L1* mRNA occurring upon *KIF18A* knockdown was in line with pro-apoptotic effect of *KIF18A* knockdown together with downregulation of anti-apoptotic *BCL2* (**Fig.S1A**). Finally, KIF18A had no effect neither on *AKT1* nor *CREB1* mRNAs expression. We have previously shown that overexpression of PUM1 strongly and PUM2 slightly upregulate apoptosis and that overexpression of both caused downregulation of TCam-2 cell cycle (Janecki et al., 2018).

### Knockdown of KIF18A caused TCam-2 cell cycle arrest in G2/M phase

Since *KIF18A* orthologue depletion or mutation in mice caused germ cell cycle arrest (Czechanski et al., 2015), we sought to check whether such an effect also takes place in human germ cell model - TCam-2. To this end, upon *KIF18A* siRNA knockdown we performed cell cycle analysis 48 h post-transfection using propidium iodide staining and flow cytometry. In these conditions we observed differences in the distribution of cell cycle phases. Namely, we found a lower number of cells in G0/G1 and S phases, in favour of cells representing G2/M phase (**Fig.5A**, upper panel). This shows that KIF18A is necessary for TCam-2 cell mitotic progression in proliferating germ cells. The efficient knockdown of KIF18A in cells compared to control cells transfected with control siRNA is illustrated on **Fig.5A** (lower panel). The quality of the separation of cells being at different cell cycle phases is shown by ModFit LT Software (Verity Software House) (**Fig.5B**).

**Fig. 5.**
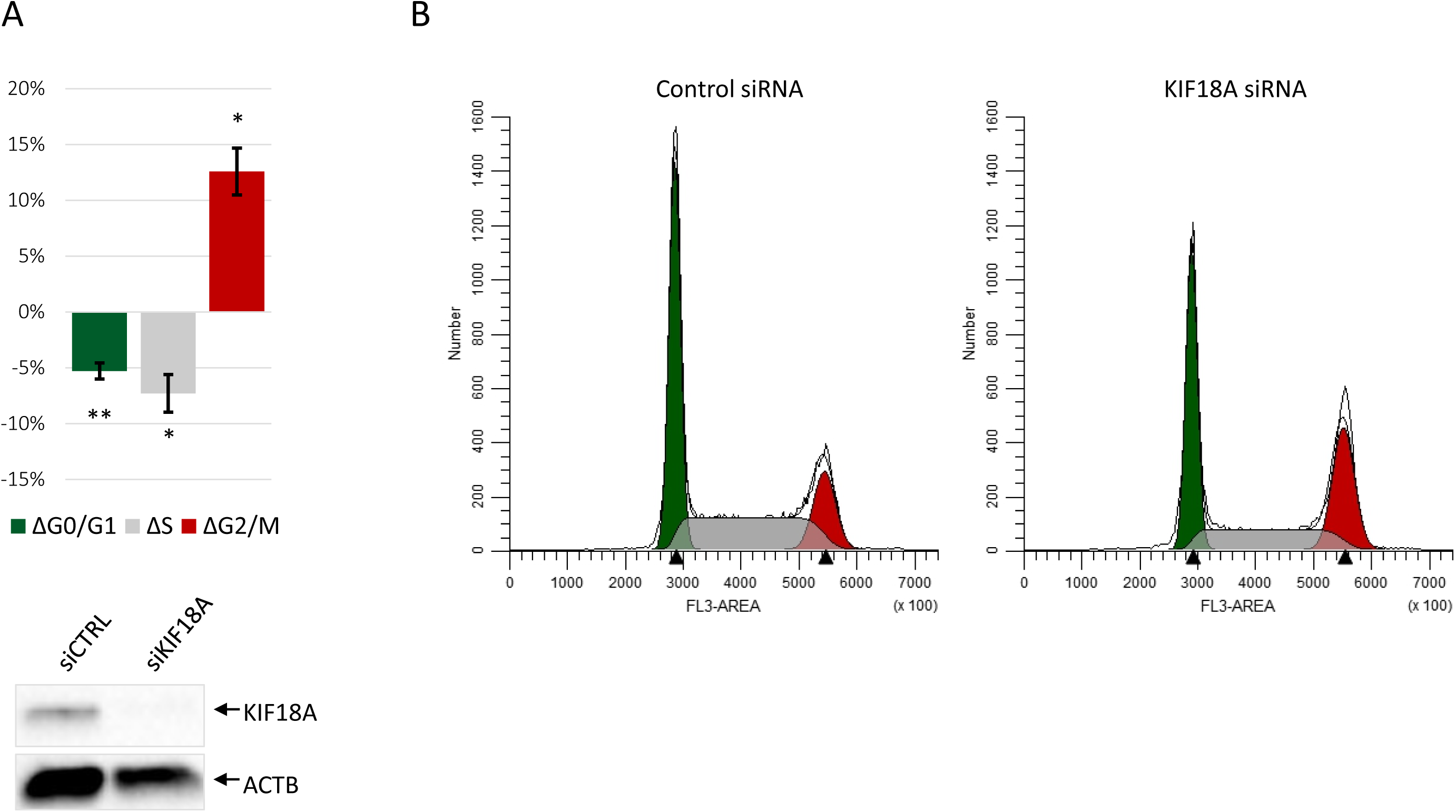
The effect of KIF18A siRNA silencing on the cell cycle, as measured by flow cytometry using propidium iodide. **A** - The cell cycle changes are represented as the % of cells in each phase (G0/G1;S; G2/M) upon KIF18A knockdown in comparison to the baseline (0%) of siRNA control (upper panel). Representative western-blot using anti-KIF18A antibody showing the level of KIF18A knockdown, compared to ACTB (anti-ACTB antibody), serving as the reference (lower panel). **B** - Representative ModFit graphs showing separation quality of particular cell cycle phases 48 h post-transfection upon control (left panel) and KIF18A siRNA silencing (right panel). All experiments were performed in 3 biological replicates with 2 technical in each. * P-value<0.05; ** P-value<0.005.

## DISCUSSION

The mechanisms responsible for regulation of expression of KIF18A, one of the kinesins playing specific roles in germ cell development, have not been investigated extensively. The presence of PBE motifs in 3’UTR suggested regulation by PUM proteins. Likewise, *KIF18A* mRNA was one of the candidates selected as PUM regulated mRNA in our recent global screens for PUM1 and PUM2 targets in TCam-2 cells (Smialek et al. [PDF] biorxiv.org, doi: https://doi.org/10.1101/760967). Here, we propose that KIF18A expression is regulated by PUM proteins for the following reasons. First, by co-immunoprecipitation we demonstrated enrichment of *KIF18A* mRNA in anti-PUM immunoprecipitates indicating interaction with PUM. Second, upon PUM siRNA silencing we observed *KIF18A* upregulation at mRNA and protein level. Third, the luciferase reporter assay upon PUM proteins overexpression or silencing shows that repression by PUM proteins is mediated by 3’UTR of *KIF18A*. Fourth, Kif18a mRNA in mice contains the same PBE UGUAHAUW motifs (not shown) in 3’UTR, indicating that this type of KIF18A posttranscriptional regulation may be evolutionarily conserved in mammals. Altogether, our results clearly show that *KIF18A* mRNA is repressed by PUMs in TCam-2 cells and that this regulation is mediated by 3’UTR. The repression by PUMs may be part of the posttranscriptional regulatory mechanism of *KIF18A* expression. To the best of our knowledge, KIF18A is the first identified mammalian functionally germ cell specific target gene that is regulated by PUM proteins.

The next important question we addressed was whether KIF18A, acting as a PUM protein target, impacts processes crucial for germ cell development such as proliferation, apoptosis and cell cycle of TCam-2 cells. If yes, whether those effects are compatible with KIF18A repression by PUM proteins. We obtained positive answers in all three contexts. Namely, the anti-proliferative effect of KIF18A depletion was in line with the effect of PUM proteins. Therefore, the activity of PUM to downregulate proliferation may be partially accounted for by repression of KIF18A. Given that PUM proteins regulate numerous RNA targets (Galgano et al., 2008) (Smialek et al. [PDF] biorxiv.org, doi: https://doi.org/10.1101/760967), there are probably other genes with similar properties to KIF18A which are worth further investigation. Subsequently, the finding that *KIF18A* depletion inhibits cell cycle in G2/M phase is consistent with previous reports showing that depletion of *KIF18A* caused mitotic arrest (Huang et al., 2009);(Liu et al., 2010). That effect is also consistent with our recent report documenting that PUM proteins downregulate TCam-2 cell cycle (Janecki et al., 2018). Therefore, we propose that posttranscriptional modulation of KIF18A level represents one of mechanisms by which PUM proteins influence the TCam-2 cell cycle. The decrease of TCam-2 cells proliferation rate upon *KIF18A* depletion is in line with cell cycle arrest in G2/M phase.

Finally, we have observed a significant pro-apoptotic effect of *KIF18A* depletion on TCam-2 cells which was in line with the recently reported effect of PUM proteins (Janecki et al., 2018). We show that this KIF18A effect might be accounted for by impacting a group of genes that are related to apoptosis. Namely, a positive influence of KIF18A depletion on pro-apoptotic *BCL2L1* gene expression as well as its negative effect on expression on anti-apoptotic *BCL2* gene expression may account for its anti-apoptotic activity in TCam-2 cells. Interestingly, we have previously reported that PUMs overexpression caused upregulation of apoptosis (Janecki et al., 2018), the same effect as KIF18A knockdown. This suggests that KIF18A is among targets of PUM proteins influencing apoptosis. Our analysis showed that KIF18A knockdown caused downregulation of anti-apoptotic *BCL2* mRNA level, which is consistent with the described anti-apoptotic role of KIF18A. However, KIF18A knockdown also caused upregulation of anti-apoptotic *CCND1* and *PIK3CA* genes which is opposite to what one could expect and therefore requires further studies.

The precise regulation of proliferation, apoptosis and cell cycle is crucial for germ cell development, and dysregulation of those processes may lead to infertility or cancer. TCam-2 cells, while representing human male germ cells, originate from a type of the most frequent type of human TGCT, the seminoma (for review see (Batool et al., 2019)). TGCTs are the most common solid tumours in young men, and their incidence is on the rise (Rosen et al., 2011). Our finding that KIF18A depletion inhibits proliferation of TCam-2 cells, is consistent with previous reports showing that KIF18A stimulated proliferation of other cancer cells *in vitro* and *in vivo*, such as clear cell renal carcinoma, breast cancer or lung adenocarcinoma (Chen et al., 2016; Zhang et al., 2010; Zhong et al., 2019). Also, the effect of KIF18A depletion, causing upregulation of apoptosis and cell cycle arrest represent features promoting cancer.

Altogether, this study broadens the current knowledge concerning regulation of *KIF18A* in germ cells. Further studies of KIF18A regulation by PUM proteins in the context of pathways governing such processes as cell proliferation, cell cycle and apoptosis is of high priority. Considering that these three processes are crucial for germ cell development and cancer, such further studies may shed light on the molecular basis of functional relationships between these two processes.

## MATERIALS & METHODS

### Cell culture and transfection

TCam-2 cells were cultured in RPMI with GlutaMAX medium (Gibco, Life Technologies) supplemented with 10% (v/v) fetal bovine serum (HyClone) and 1% (v/v) antibiotic antimycotic solution (Lonza). The cells were transfected with plasmid (pCMV6-entry from Origene) and siRNA constructs (Santa Cruz Biotechnology) using the Neon Transfection System (Life Technologies) according to the manufacturer’s protocol.

### Luciferase assays

For luciferase assays, 2×10^5^ TCam-2 cells were transfected with 1.5 μg of plasmid DNA encoding PUM1, PUM2, or an empty plasmid in the pCMV6-entry vector system (OriGene Technologies), plus the full-length 3□UTR of *KIF18A* in the psiCheck2 dual luciferase vector system (Promega, Germany) in 10:1 (plasmid DNA:luciferase vector) ratio. Transfected cells were cultured in 12-well plates for 24 h in standard medium without antibiotic antimycotic solution. For *PUM* transient knockdown experiments, 1.5×10^5^ TCam-2 cells were transfected with 40 nM siRNA and 150 ng of psiCheck2 vector constructs as described above, and then cultured in 12-well plates for 48 h to achieve effective *PUM* knockdown. Transfections were performed in three biological repetitions. Cells were lysed and luminescence from each well was measured twice using a Glomax-Multi Detection System luminometer (Promega) and the Dual-Luciferase Reporter Assay (Promega) according to the manufacturer’s protocol. Average Renilla to firefly luciferase luminescence ratios and standard deviations were calculated from three experiments. Luminescence ratios for each combination of constructs and/or siRNA were presented as a percent of relative luciferase units (RLU). The sample transfected with empty pCMV6-entry or control siRNA plus the reporter construct was considered as 100%.

### MTS proliferation assay

TCam-2 cells were transfected with 40 nM siRNA mixture of 3 different *KIF18A* sequences (Santa Cruz Biotechnology, sc-96629), *PUM1* (Santa Cruz Biotechnology, sc-62912), *PUM2* (Santa Cruz Biotechnology, sc-44773) and control scrambled siRNA (Santa Cruz Biotechnology, sc-37007) (sequences in **Table S1**). These were done in three biological replicates (three independent transfections; 7 technical repeats/wells on a 96-well plate) and were cultured for 24, 48, 72, 96 and 120 h after transfection. To obtain a similar number of viable cells per well the same (5,000 cells per well) number of cells were seeded to 96-well plates directly after transfection. The cell viability was measured using GLOMAX (Promega) at 450 nm (and 750 nm for cell background) wavelength 24, 48, 72, 96 and 120 h after transfection and 4 h after adding 20 µl of CellTiter 96® AQ One Solution Reagent (Promega) to each well containing transfected TCam-2 cells.

The expression of KIF18A, PUM1, PUM2 and ACTB proteins was measured 24, 48, 72, 96 and 120 h after transfection by western blot using anti-KIF18A, -PUM1, -PUM2 and -ACTB antibodies. ACTB expression was used as a reference to estimate the protein content in each blot.

### Western blotting

Overexpression and silencing efficiency was measured by western blot analysis under standard conditions, using a nitrocellulose membrane, horseradish peroxidase (HRP)-conjugated secondary antibodies, and semi-quantitative measurement of protein levels was performed using ImageLab 5.1 software (BioRad). The chemiluminescent signal was detected using Clarity™ Western ECL Substrate for HRP (BioRad).

### Antibodies

For the measurements of knockdown efficiency, the primary antibody anti-KIF18A (Thermo Fisher Scientific, PA5-31477, 1:5000), anti-PUM1 (Santa Cruz Biotechnology, sc-65188, 1:1000) and anti-PUM2 (Santa Cruz Biotechnology, sc-31535, 1:250) were used. The primary antibody anti-DDK (FLAG^®^) (OriGene Technologies, TA50011, 1:1000) was used for detection of proteins overexpressed from the pCMV6-entry vector. Anti-ACTB reference protein (Sigma Aldrich, A2066, 1:10000) antibody was also used. Goat anti-rabbit IgG-HRP (Sigma Aldrich, A6154, 1:25000), mouse anti-goat IgG-HRP (Santa Cruz Biotechnology, sc-2354, 1:50000) and mouse IgGκ-HRP (Santa Cruz Biotechnology, sc-516102, 1:10000) were used in the study as a secondary antibodies.

### Flow cytometry

Cell cycle analysis was performed 48 h after TCam-2 transfection with 40 nM control and KIF18A siRNAs. For this purpose, TCam-2 cells were washed with PBS and fixed in cold 100% methanol on ice for 10 minutes. After incubation at 37°C for 15 minutes in 50 µg/ml propidium iodide (PI) (Sigma Aldrich) containing 330 µg/ml RNase A (Sigma Aldrich), cells were incubated for 1 h on ice and finally measured using the S3e™ Cell Sorter (BioRad) apparatus. Data files were analysed using ModFit LT (Verity Software House).

Detection of apoptotic TCam-2 cells was performed 48 h after transfection of TCam-2 cells with control and KIF18A siRNA constructs using an Annexin V-FITC Apoptosis Detection Kit (Beckman Coulter), according to the manufacturer’s protocol, followed by flow cytometry using a FlowSight instrument (Amnis). The results were analysed using Image Data Exploration and Analysis Software version 6.0 (IDEAS^®^ v6.0, Amnis). For all flow cytometry experiment 3 biological repetitions with 2 technical repetitions in each were performed.

### RNA co-immunoprecipitation

For RNA co-immunoprecipitation experiments 2×10^6^ TCam-2 cells were transfected with 15 µg of pCMV6-entry empty (only DDK), and PUM1-DDK and PUM2-DDK coding sequences. 24 h post-transfection TCam-2 cells were washed twice with ice-cold PBS and subjected to UV cross-linking at 254 nm on a HEROLAB CL-1 Cross-linker for 60 s (0.03 J). For one RIP reaction, 2-3×10^6^ cells were lysed in 1000 μl of RIPA Lysis Buffer (with 1% Triton X-100) for 30 min with rotation at 4 °C. Lysates were centrifuged, and the supernatant was mixed with Anti-FLAG M2 magnetic beads (Sigma Aldrich, M8823, 50µl) with protease and RNase inhibitors. The RIP reaction was held for 3 h at 4 °C on a rotator in a final volume of 1 ml. Then, magnetic beads were washed three times with RIPA buffer (without Triton X-100) followed by treatment with proteinase K at 55° C for 30 minutes. Total RNA was isolated from 80% magnetic beads using a TRIzol® Reagent (Life Technologies) according to the manufacturer’s protocol. 20% of the magnetic beads, 3% of input and flow-through were loaded on 8% PAGE gels for western blot analysis.

### Real-Time quantitative PCR

To measure KIF18A mRNA enrichment in RIP anti-PUM1 and PUM2, to the total RNA from magnetic beads after RNA-co-immunoprecipitation (50-100 ng) were added equally 200 ng of *D. melanogaster* total RNA (spike-in RNA) for later reverse transcription and RT-qPCR experiments (in which Act5c mRNA from *D. melanogaster* was treated as reference).

To measure *KIF18A, PUM1, PUM2, AKT1, BCL2, BCL2L1, CREB1, CCND1* and *PIK3CA* mRNA levels, total RNA from cell cultures was isolated using TRIzol® Reagent (Life Technologies) according to the manufacturer’s protocol. RNA was treated with DNase I (Sigma Aldrich) and reverse-transcribed using the Maxima First Strand cDNA Synthesis Kit (Life Technologies) according to the manufacturer’s protocol. Total cDNA was used as a template for qPCR amplification. The reaction was carried out using the CFX96 Touch™ Real-Time PCR Detection System (Bio-Rad, Poland) in 10 μl volumes containing 10 mM Tris-HCl, pH 8.3, 50 mM KCl, 3.5 mM MgCl_2_, 0.2× Sybr Green, 0.2 mM dNTP mix (dATP, dCTP, dGTP, dTTP), 0.2 μM F and R primers and 0.5 U JumpStart™ Taq DNA Polymerase (Sigma Aldrich, D4184). Specific primers are listed in **Table S2**. Amplification parameters were as follows: initial denaturation 95° C, 2.5 min and 40 cycles of: denaturation 95° C, 15 sec, annealing 10 sec (annealing temperatures for each primer pair are given in **Table S2**), extension 72° C, 15 sec. ACTB and GAPDH were used for normalisation.

### Statistical analysis

One-way unpaired t-test was used to estimate statistical significance. A *P-value* < 0.05 (*) was considered statistically significant.

## ACKNOWLEDGMENTS

We thank Anna Spik for excellent technical assistance,Jaruzelska lab and Simon Anthony Patterson for reading and correcting the manuscript. This study was supported by grants from the National Science Centre Poland no 2013/09/B/NZ1/01878 to JJ and ETIUDA6 scholarship no 2018/28/T/NZ1/00015 to MJS.

## CONFLICT OF INTERESTS

The authors declare no conflict of interest.

---

**Fig.S1** Influence of KIF18A on some mRNAs encoding anti- or pro-apoptotic proteins. **A** - RT-qPCR results of anti-apoptotic *AKT1, BCL2, CREB1, CCND1* and *PIK3CA* and pro-apoptotic shorter isoform of *BCL2L1* mRNA expression level changes upon KIF18A depletion (48 and 72 h post-transfection) in comparison to control siRNA (siCTRL) and in reference to ACTB and GAPDH expression level. **B** - Representative western-blot showing the level of KIF18A siRNA knockdown 48 and 72 h post-transfection, in comparison to control siRNA, ACTB detected using anti-ACTB antibody served as reference. All experiments were performed in 3 biological replicates with 5 technical in each. *** P-value<0.0005; **** P-value<0.00005

